# Identification of Evolutionarily Conserved VSX2 Enhancers in Retinal Development

**DOI:** 10.1101/2023.10.17.562742

**Authors:** Victoria Honnell, Shannon Sweeney, Jackie Norrie, Cody Ramirez, Beisi Xu, Brett Teubner, Ah Young Lee, Claire Bell, Michael A. Dyer

## Abstract

Super-enhancers (SEs) are expansive regions of genomic DNA that regulate the expression of genes involved in cell identity and cell fate. Recently, we found that distinct modules within a murine SE regulate gene expression of master regulatory transcription factor *Vsx2* in a developmental stage- and cell-type specific manner. *Vsx2* is expressed in retinal progenitor cells as well as differentiated bipolar neurons and Müller glia. Mutations in *VSX2* in humans and mice lead to microphthalmia due to a defect in retinal progenitor cell proliferation. Deletion of a single module within the *Vsx2* SE leads to microphthalmia. Deletion of a separate module within the SE leads to a complete loss of bipolar neurons, yet the remainder of the retina develops normally. Furthermore, the *Vsx2* SE is evolutionarily conserved in vertebrates, suggesting that these modules are important for retinal development across species. In the present study, we examine the ability of these modules to drive retinal development between species. By inserting the human build of one *Vsx2* SE module into a mouse with microphthalmia, eye size was rescued. To understand the implications of these SE modules in a model of human development, we generated human retinal organoids. Deleting one module results in small organoids, recapitulating the small-eyed phenotype of mice with microphthalmia, while deletion of the other module leads to a complete loss of ON cone bipolar neurons. This prototypical SE serves as a model for uncoupling developmental stage- and cell-type specific effects of neurogenic transcription factors with complex expression patterns. Moreover, by elucidating the gene regulatory mechanisms, we can begin to examine how dysregulation of these mechanisms contributes to phenotypic diversity and disease.

**Summary Statement:** Herein, we describe how conserved modules within a single super-enhancer can regulate *VSX2* gene expression across species in both mice and human retinal organoids.

## Introduction

For proper retinal development to occur, thousands of genes must be turned on and turned off in a precise spatiotemporal order (Livesey and Cepko, 2001). Many genes encoding master regulatory transcription factors are expressed in retinal progenitor cells during early stages of development and in a subset of differentiated cells at late stages of development (Haubst et al., 2004, (Castro et al., 2011, (Nishida et al., 2003). For example, *Sox2* is expressed in retinal progenitor cells and persists in differentiated Müller glia in the adult retina (Graham et al., 2003). Likewise, *Pax6* is expressed in retinal progenitor cells at early stages of development and persists in amacrine cells and Müller glia (Collinson et al., 2003, (Marquardt et al., 2001).

While these complex cell type-specific gene expression patterns have been well-characterized during retinogenesis, the underlying molecular mechanisms by which these genes are regulated is not well understood.

Prior studies have computationally identified hundreds of putative super-enhancers across multiple stages of retinal development (Aldiri et al., 2017, (Marchal et al., 2022). A major challenge in the field is correctly identifying enhancers and their target genes in a developmental stage and cell-type specific manner. We recently identified a super-enhancer (SE) upstream of the *Vsx2* gene that is necessary and sufficient for the complex expression pattern of *Vsx2* during development (Honnell et al., 2022). *Vsx2* is expressed in retinal progenitor cells and maintained in differentiated bipolar neurons and Müller glia (Liu et al., 1994, (Rowan and Cepko, 2004, (Vitorino et al., 2009, (Buenaventura et al., 2018). *Vsx2* gene knockout mice (*OrJ*) are born with microphthalmia, a very small eye, due to a defect in retinal progenitor cell proliferation (Burmeister et al., 1996, (Ferda Percin et al., 2000, (Livne-Bar et al., 2006, (Truslove, 1962).

Thus, determining the precise role of *Vsx2* in differentiated bipolar neurons and Müller glia in this model is difficult because normal retinal development does not occur.

We found that, in mice, the developmental stage-specific and cell-type specific expression pattern of *Vsx2* is achieved by distinct enhancer modules within its adjacent SE (Honnell et al., 2022). One module (Region 0) is responsible for retinal progenitor cell proliferation. Deletion of this region leads to microphthalmia. A separate module (Region 3) regulates bipolar cell genesis. Deletion of this region leads to complete loss of a single cell type, the bipolar neurons. Bipolar neurons are present in the Region 0 deletion retina and the eye is of a normal size in the Region 3 deletion retina demonstrating that the *Vsx2* gene is regulated by a modular SE. Thus, by knocking out the enhancers cis to the *Vsx2* gene, we are able to control *Vsx2* gene expression without perturbing the gene itself.

Considering that these sequences are evolutionarily conserved across vertebrates, we hypothesize that they are necessary for normal human retinal development and interchangeable between species. In the present study, we discovered that the human VSX2 enhancer modules are sufficient for driving reporter expression in the mouse retina. We then assessed the ability of the R0 human enhancer to function between species by replacing the mouse enhancer with the human enhancer in vivo. Using the Region 0 deletion mouse, which has microphthalmia, we knocked-in the human Region 0 enhancer and found that eye size and vision were rescued. To understand how these enhancers may function in a model of human retinal development, we generated Region 0 and Region 3 knockout retinal organoids from human embryonic stem cells (hESCs). These data suggests that the *Vsx2* enhancer elements are functionally interchangeable between humans and mice, and that both enhancers are necessary for normal human retinal organoid development. This work further explores enhancer mechanisms regulating *Vsx2* gene expression and could offer insights for previously unknown enhancer candidates contributing to microphthalmia and blindness in humans.

## Results

### The *Vsx2* SE enhancer regions are evolutionarily conserved

Using integrated epigenetic analyses, we previously identified a 41kb SE upstream of the *Vsx2* gene in mice (Norrie et al., 2019). Within the *Vsx2* super-enhancer are four evolutionarily conserved regions, R0-37, R1-28, R2-22, and R3-17 located 37, 28, 22, and 17 kilobases from the *Vsx2* mouse promoter, respectively (Honnell et al., 2022) (Fig. S1A). To determine whether these regions are sufficient for driving reporter expression in mice, we subcloned them into a pP reporter plasmid driven by a minimal promoter, as previously described (Kim et al., 2008). Mouse Regions 0-3 were individually co-electroporated into mouse retina at P0 in vivo with a constitutive H3.3 Scarlet reporter plasmid, labeling all electroporated cells. We scored the proportion of GFP+, Scarlet+/Scarlet+ cells for each cell type for each reporter plasmid. R0-37 is sufficient for driving reporter expression in Müller glia and R3-17 is sufficient for driving reporter expression in bipolar neurons (Fig. S1B). Since the *Vsx2* SE extends into the neighboring *Lin52* gene, we also examined evolutionarily conserved regions within this gene. However, no statistically significant reporter expression was observed (Fig. S1C). Considering that conserved Regions 0-3 are several kilobases in size, we refined them to sequences conserved only in distantly related species. Upon refining Region 3, we found that Region 3-d exhibited robust transgene expression in the bipolar neurons compared to Region 3-c (Fig. S1D). Notably, Region 3-d contains a 164 bp element that was previously found to drive reporter expression in rat bipolar neurons by the Cepko laboratory (Kim et al., 2008). Lastly, we combined each of the refined regions to create a “mini-enhancer” and found that it recapitulates the cell-type specific expression inherent to its individual elements (Fig. S1E).

Considering that there is strong evolutionary conservation of these genomic sequences across vertebrates, we then examined the ability of these regions to drive reporter expression between species. To determine if the human genomic sequences of Regions 0-3 (namely hR0-36, hR1-25, hR2-20, and hR3-16) are sufficient for driving reporter expression in mice, we subcloned them into the reporter plasmid described above, and electroporated them into mouse retina (Fig. 1A-B). When electroporated into a mouse, hR0-36 is sufficient for driving reporter expression in Müller glia and hR3-16 is sufficient for driving reporter expression in bipolar neurons (Fig. 1C-D). This is consistent with the results observed from electroporation of the ortholog mouse enhancer sequences (Fig. 1E). Taken together, these results suggest that the *VSX2* SE contains evolutionarily conserved enhancer elements that are sufficient for driving cell-type specific reporter expression between species.

**Figure 1.**
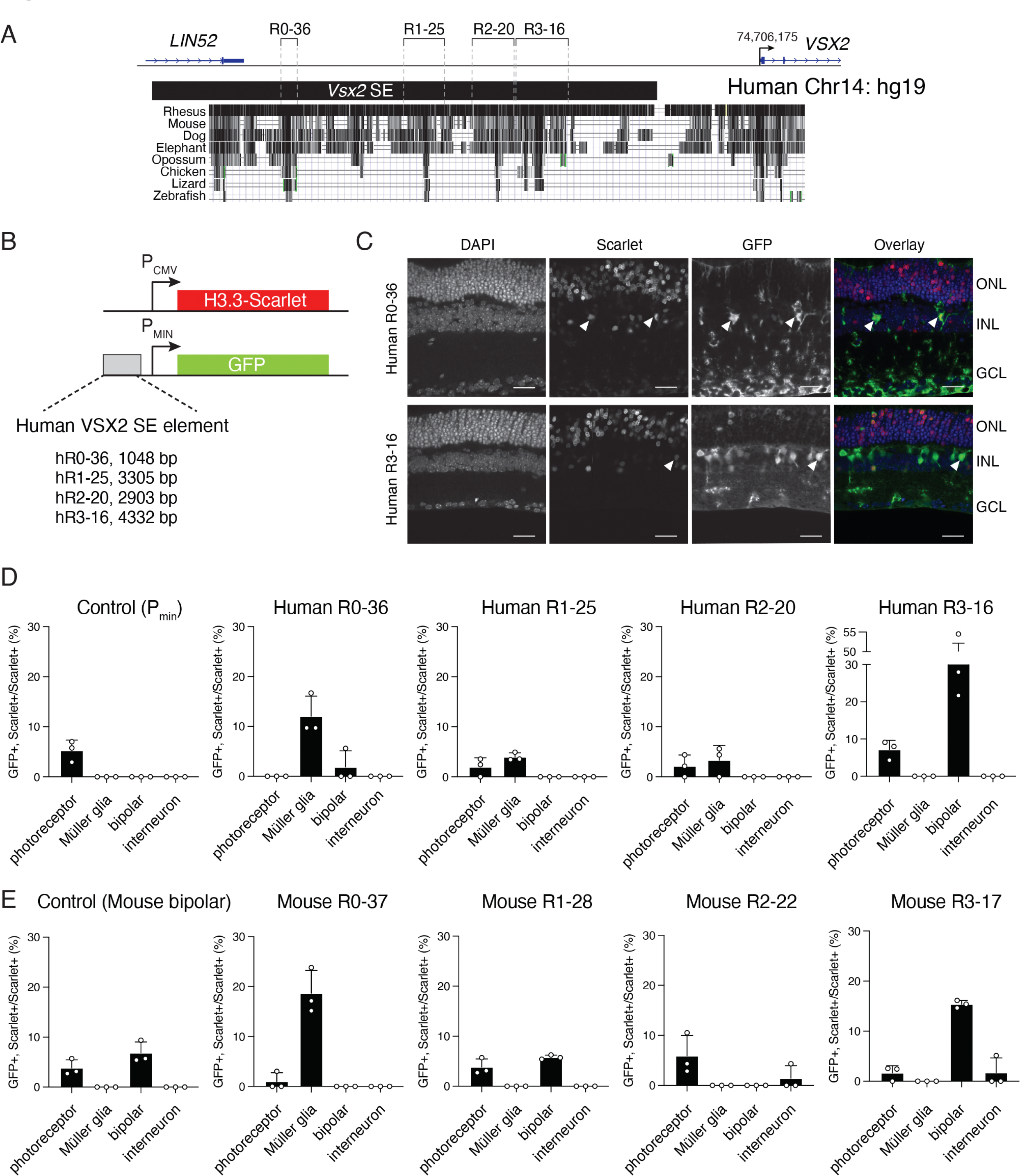
The VSX2- SE contains evolutionarily conserved enhancer elements that are sufficient for driving reporter expression between species. A Drawing of the original *Vsx2* SE identified by H3K27Ac ChIP-seq (black bar) and the original deletions with coordinates in mm10. Evolutionarily conserved sequences across vertebrates are displayed below each Region deletion (black segments). B Drawing of plasmids used for reporter assays for in vivo square wave electroporation at P0 in mice and harvested at P21. A minimal promoter (P_MIN_) that is not sufficient for high-level expression is upstream of GFP and a strong constitutive promoter (P_CMV_) is used for the Scarlet reporter. C Micrographs of GFP (green) and Scarlet (red) expression at P21 from square wave electroporation of P0 mice. Arrows indicate Müller glia for the R0 fragment and bipolar neurons for the R3 fragment. Scale bar: 25 uM. D Bar plot showing mean and standard deviation of three biological replicates for each human reporter construct. The negative control is the minimal promoter without a subcloned Region and has very little expression E Bar plot showing mean and standard deviation of three biological replicates for each mouse reporter construct. The positive control has a previously identified bipolar-specific element. Abbreviations: ONL, outer nuclear layer; INL, inner nuclear layer; GCL, ganglion cell layer.

### The *Vsx2* SE Region 0 is functional between species

Mutations in *VSX2* lead to microphthalmia in humans and mice (*OrJ)* due to a defect in retinal progenitor cell proliferation (Burmeister et al., 1996, (Livne-Bar et al., 2006, (Ferda Percin et al., 2000, (Truslove, 1962). Our previous work found that microphthalmia also occurs in Region 0 deletion mice (R0-37*^Δ/Δ^*) (Honnell et al., 2022) (Fig. 2A-C). While the *OrJ* and R0- 37*^Δ/Δ^*mice are blind, their ability to photoentrain has not been explored (Honnell et al., 2022). We examined the ability of *OrJ* and R0-37*^Δ/Δ^* mice to photoentrain by assessing running-wheel activity accompanied by light schedule shifts over 36 days. Despite having photoreceptors, melanopsin, and an optic nerve, the *OrJ* and R0-37*^Δ/Δ^* mice do not photoentrain (Fig. S2 A-E).

**Figure 2.**
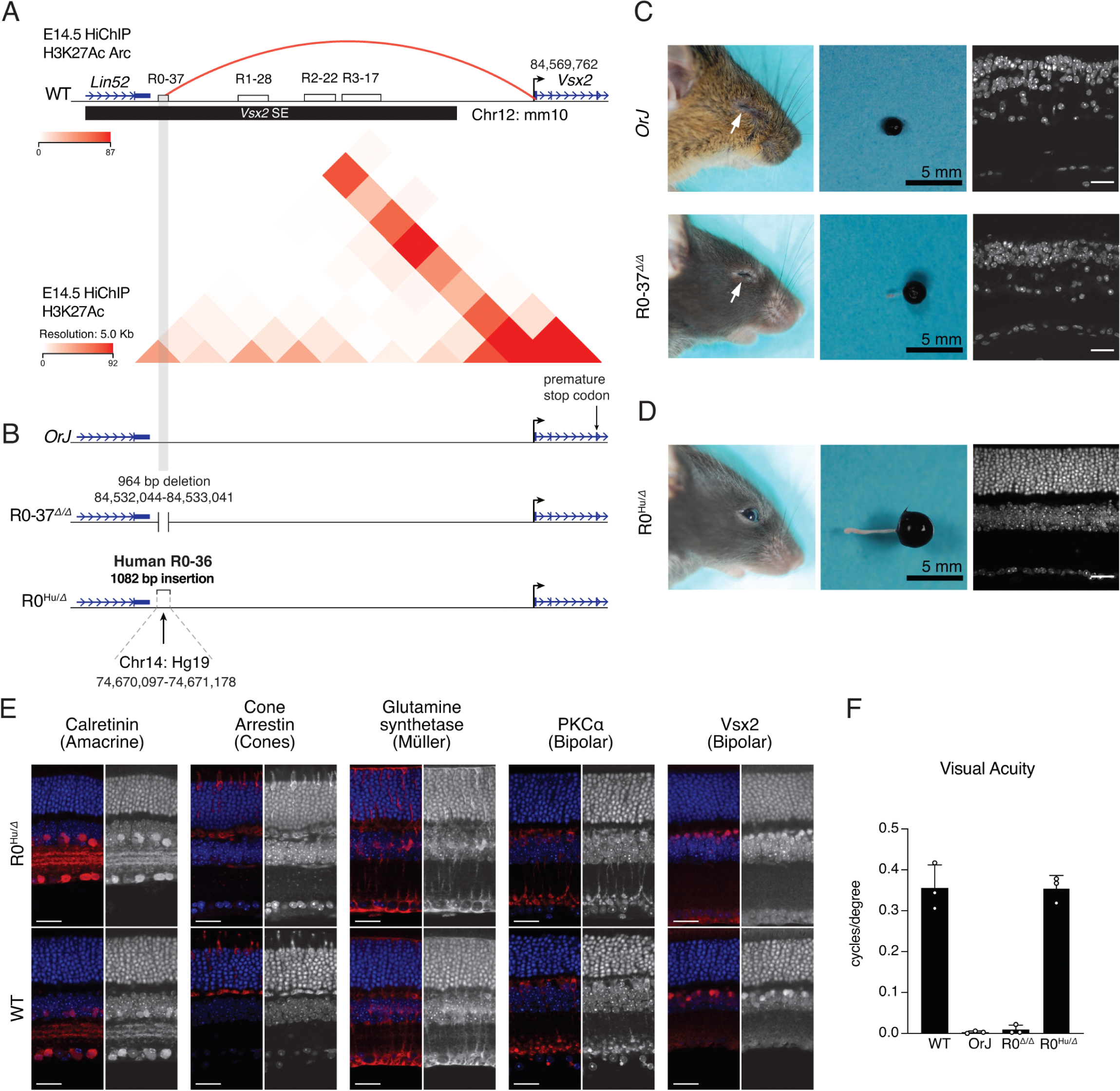
Human enhancer rescues eye size and vision in the R0 deletion mouse. A. Drawing of the *Vsx2* SE (black bar) containing evolutionarily conserved Regions 0-3 in mm10. There is a significant chromatin interaction between R0-37 and the *Vsx2* promoter (red arc) as determined by integration of HiChIP and ChIP-seq. H3K27ac data. Heat map displaying chromatin interactions within the *Vsx2* SE of E14.5 retina determined by HiChIP (below). **B** The three mouse lines examined and their respective mutations. **C** Photograph of eyes and micrograph of retina for *orJ* and R0-37^Δ/Δ^ adult mice as a reference for microphthalmia. **D** Photograph of eyes and micrograph of retina for R0^Hu/Δ^ mice. **E** Images of retinal cell types at 40X. **F** Bar plot of mean with standard deviation of photopic vision (cycles/degree). n= 3 biologically independent mice for each mouse strain. Scale bars: 25 uM.

Considering that R0-37 is necessary for normal eye size, we next examined how this enhancer region may promote RPC proliferation during early periods of retinal development. By integrating HiChIP and ChIP-seq. data for mark H3K27ac, we found a significant chromatin interaction between R0-37 and the *Vsx2* promoter in E14.5 wildtype retina (Fig. 2A).

To examine functional capabilities of the Human R0 enhancer between species, we knocked-in the Human R0 enhancer, hR0-36, into the R0-37 deletion mouse using CRISPR- Cas9, thereby replacing the mouse enhancer sequence with the human ortholog (Fig. 2B). The Human Region 0 insertion mouse (R0*^Hu/Δ^*) displays a normal eye size as well as all three nuclei layers as determined by morphological analysis (Fig. 2D). The R0*^Hu/Δ^* retina also contains a normal distribution of rod photoreceptors, cones, bipolar and amacrine neurons, and retinal ganglion cells as observed by immunofluorescence (Fig. 2E). Furthermore, these mice have normal visual acuity as determined by optometry (Fig. 2F). These datasets suggest that hR0-36 is able to rescue mouse eye size and restore visual acuity, and is thus functional across murine retinal development.

### Vsx2 SE Region 0 is necessary for normal human retinal organoid development

To understand the effect of the *Vsx2* SE in a model of human retinal development, we generated human retinal organoids following an established protocol with three defined developmental stages (Capowski et al., 2019) (Fig 3A). Using CRISPR-Cas9, we deleted hR0-36 (R0-36*^Δ/Δ^*), hR3-16 (R3-16*^Δ/Δ^*), and generated frameshift mutations in exon 2 for a VSX2 null (VSX2^-/-^) in human embryonic stem cells (hESCs) (Fig. 3B). Two sublines were created and analyzed for each deletion. No differences in sublines were observed by RNA seq, qRT-PCR, or morphological analysis by IHC (Supplemental Dataset 1). The parent cell line contains a dual reporter that labels cells expressing *VSX2* and *CRX* (H9 *VSX2*-GFP/*CRX*-TdTomato) (Fig. 3C). Retinal organoids were collected for analysis across the three developmental stages (Capowski et al., 2019). At Stage 1, all retinal organoid lines express *CRX*-Tomato, marking photoreceptors.

**Figure 3.**
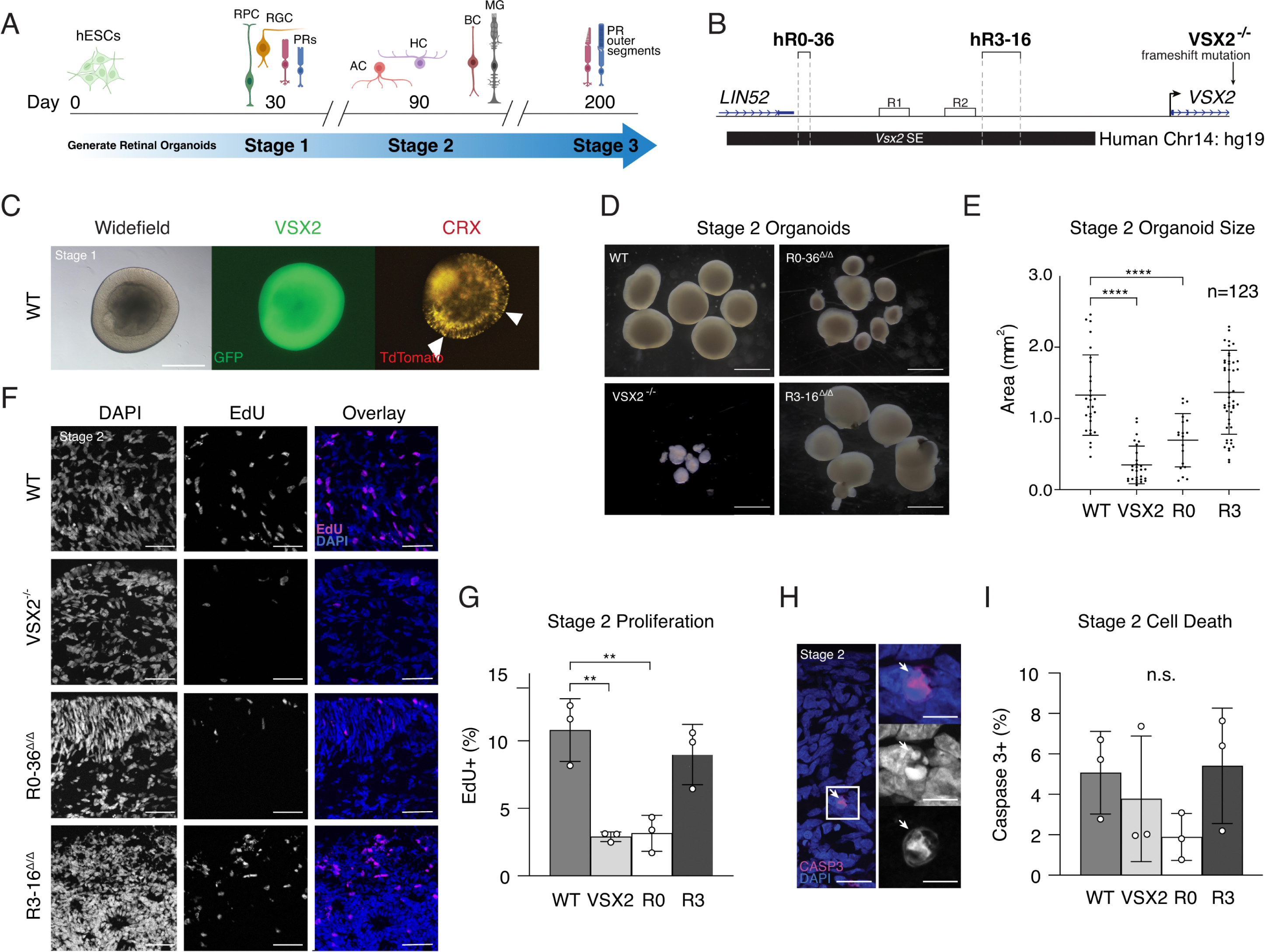
Region 0 of the VSX2-SE is necessary for normal human retinal organoid development. A. Timeline of retinal organoid generation. Stage 1 organoids contain retinal progenitor cells (RPC), retinal ganglion cells (RGC), and photoreceptors (PR). Stage 2 organoids contain amacrine cells (AC), horizontal cells (HC), and bipolar cells (BC). Stage 3 organoids contain Müler glia (MG) and developed PR outer segments. **B** Drawing of the VSX2 SE (black bar) containing evolutionarily conserved Regions 0-3 in genome build hg19. hR0-36 (R0*^Δ/Δ^*) and hR0-36 (R3^Δ/Δ^) were deleted using CRISPR-Cas9 in hESCs (location in bold). A VSX2 knock- out cell line was generated by creating frameshift mutations in the gene. **C** Stage 1 photographs of wildtype organoids taken on a widefield microscope. GFP expression is observed in WT and R3^Δ/Δ^ organoids. TdTomato is expressed across organoid lines. Scale bar: 0.5 mm. **D** Stage 2 photographs of retinal organoid lines. The core has darkened for most lines, indicative of Stage 2. **E** Scatter dot plot of organoid area in mm^2^. Each dot represents a single retinal organoid. VSX2^-/-^ and R0^Δ/Δ^ retinal organoids are significantly smaller than wildtype retinal organoids (unpaired two-tailed t-test, **** p < 0.0001). There is no significant difference observed between R3^Δ/Δ^ and WT retinal organoid area (unpaired two-tailed t-test, p = 0.7811). **F** Micrograph of retinal organoids from each line showing EdU labeling (magenta) and DAPI (blue). **G** Bar plot showing the mean and standard deviation of the proportion of EdU+ cells scored from micrographs of retinal organoid sections. VSX2^-/-^ and R0-36^Δ/Δ^ retinal organoids display a significant reduction in proliferation compared wildtype retinal organoids (unpaired two-tailed t- test, ** p < 0.008). **H** Micrograph of DAPI (blue) and activated caspase three (magenta) immunostaining of Stage 2 sections from a WT retinal organoid. Scale bar: 10 uM. **I** A bar plot showing the scoring for the proportion of caspase three across retinal organoid lines (n.s., not significant). Mean and standard deviation are shown.

Wildtype and R3-16*^Δ/Δ^* retinal organoids expressed *VSX2*-GFP, marking retinal progenitor cells. GFP was not visible by fluorescent light microscope in VSX2^-/-^ and hR0-36*^Δ/Δ^* retinal organoids (data not shown).

By the end of Stage 2, the majority of retinal progenitor cells are expected to have fully differentiated, and all neuronal cell types are present (Capowski et al., 2019). We observed that Vsx2^-/-^ and hR0-36*^Δ/Δ^*retinal organoids are significantly smaller compared to WT retinal organoids, as assessed by measurements of 123 traced retinal organoids from photographs (Fig. 3D-E). To determine if there is any perturbation in retinal progenitor cell proliferation as a result of the deletions, we performed EdU labeling during Stage 2 on day 90 . Retinal organoids received EdU and 1 hour later they were harvested, stained for EdU and DAPI, and scored.

There is no significant difference in size or proportion of proliferating cells between WT and R3*^Δ/Δ^* retinal organoids. However, there is a significant reduction in the proportion of EdU+ cells in the outer layer of VSX2^-/-^ and R0*^Δ/Δ^*retinal organoids compared to WT retinal organoids (Fig. 3F-G). To determine whether discrepancies in retinal organoid size was due to apoptotic retinal neurons, we performed immunostaining for activated Caspase 3 and scored the proportion of immunopositive cells. There was no significant difference in the proportion of outer layer apoptotic cells between retinal organoid lines (Fig. 3H-I). Taken together, our data suggests that the hR0-36 enhancer is necessary for retinal progenitor cell proliferation and that the R0-36*^Δ/Δ^* human retinal organoids recapitulate the small-eye phenotype observed in mice with microphthalmia.

### R0*^Δ/Δ^* human retinal organoids have aberrant gene expression across developmental stages

During early development at Stage 1, there are no visible differences between R0-36*^Δ/Δ^* and WT retinal organoids (Fig. S3 A-B). To examine potential differences in gene expression, we harvested multiple biological replicates for each organoid line across the three developmental stages for bulk RNA-seq and qRT-PCR. At Stage 1 and Stage 2, principal component analysis (PCA) plots display R0-36*^Δ/Δ^* clustered with VSX2^-/-^ retinal organoids, and R3-16*^Δ/Δ^* clustered with WT organoids, suggestive of similarity in gene expression profiles (Fig. 4A-B). We then compared R0-36*^Δ/Δ^* to Vsx2^-/-^ retinal organoids, and found that the transcriptomic data is positively correlated across stages (Fig. 4C-D).

**Figure 4.**
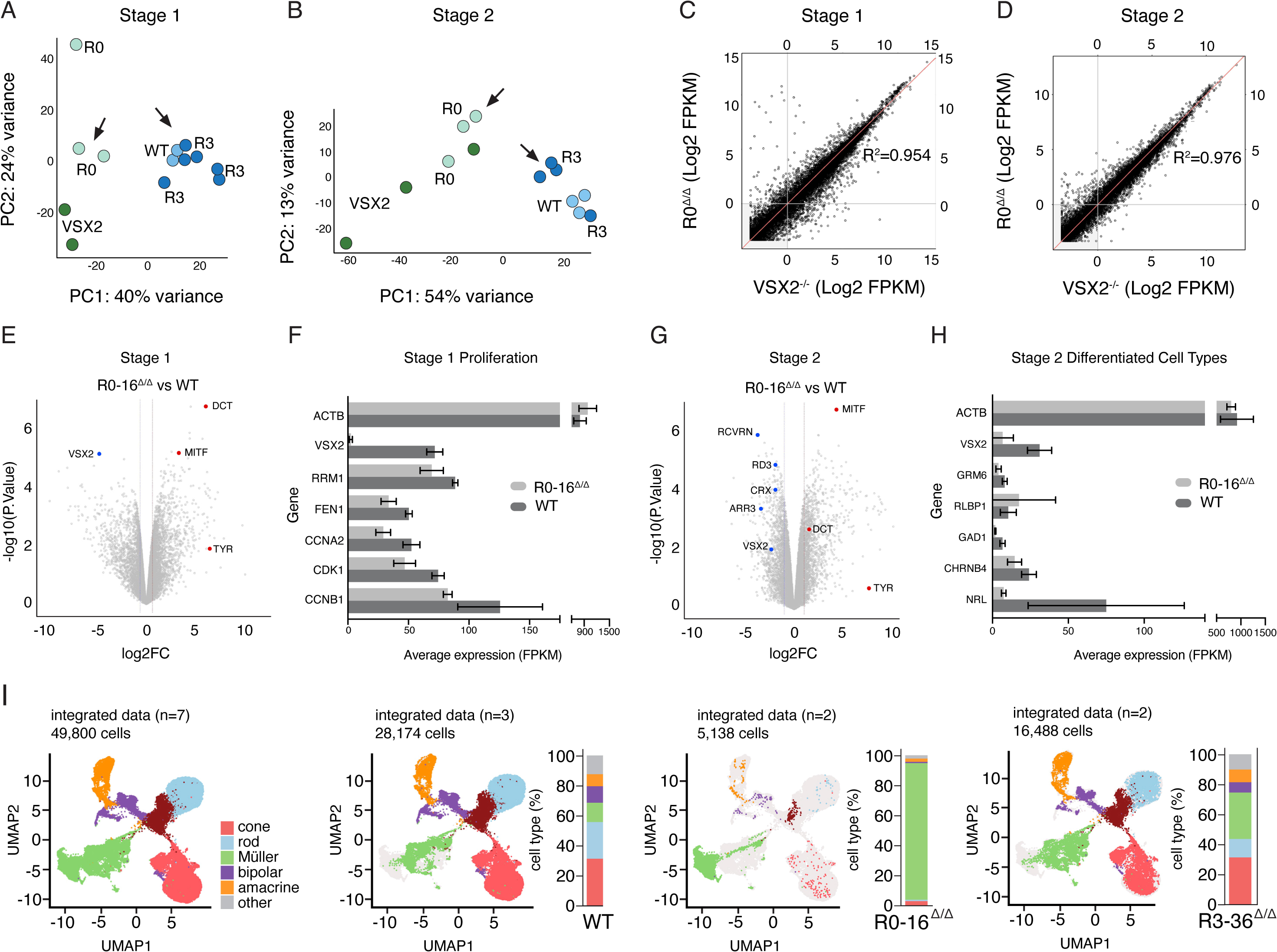
Region 0 is necessary for normal human retinal organoid development. A,B. Stage 1 and 2 principal component analysis (PCA) of bulk RNA-seq for each retinal organoid line. R3- 16^Δ/Δ^ and WT retinal organoids cluster together on the right. R0-36^Δ/Δ^ and VSX2^-/-^ retinal organoids cluster to the left. **C, D** Scatterplot displaying genes from RNA-seq. Stage 1 retinal organoids have a correlation coefficient of 0.954. Stage 2 retinal organoids have a correlation coefficient of 0.976. Two biological replicates from each line were used. **E, G** Volcano plot showing differentially expressed genes of R0-16^Δ/Δ^ vs WT Stage 1 or Stage 2 retinal organoids. Downregulated genes are shown in blue. Upregulated genes are shown in red. Two biological replicates were used for each organoid line, except for Stage 2 WT which consists of three biological replicates. **F**, **H** Bar plot of the average FPKM at Stage 1 and Stage 2 for cell cycle or retinal differentiation genes, respectively. **I** UMAPs displaying cell types and their proportions of Stage 3 human retinal organoids for all samples (left). Displayed are three WT (middle left), two R0-36^Δ/Δ^ (middle right), and two R3-16^Δ/Δ^ (right) samples.

Differential gene expression of R0-36*^Δ/Δ^* and WT retinal organoids was then examined across both stages. At Stage 1, R0-36*^Δ/Δ^*retinal organoids display downregulated *VSX2* and upregulated *Microphthalmia-associated Transcription Factor (MITF),* a target of VSX2- mediated gene repression (Horsford et al., 2005, (Rowan et al., 2004, (Bharti et al., 2008, (Zou and Levine, 2012)(Fig. 4E). This data aligns with a previous study that found that human retinal organoids derived from a patient with *VSX2*-mediated microphthalmia also exhibit upregulation in *MITF* compared to a non-affected sibling (Phillips et al., 2014). Additional upregulated genes include targets of *MITF,* namely *DCT* and *TYR (Kawakami and Fisher, 2017)* (Fig. 4E).

Considering that the R0-36*^Δ/Δ^* retinal organoids are smaller in size, display a defect in proliferation, and mirror the gene expression profile observed in VSX2^-/-^ organoids known to recapitulate microphthalmia in humans, we then asked whether cell cycle genes are also affected. Using four R0-36*^Δ/Δ^* and four WT biological replicates, we examined genes involved in cell cycle and proliferation. Overall, we observed a reduction in the average FPKM values of cell cycle genes in R0-36*^Δ/Δ^*retinal organoids compared to WT (Fig. 4F).

By Stage 2, R0-36*^Δ/Δ^* retinal organoids exhibit downregulated photoreceptor genes including cone genes *ARR3, RD3, RCVRN*, and *CRX* (Fig. 4F). *MITF* and its targets, *DCT* and *TYR,* remained upregulated (Fig. 4G). Using three R0-36*^Δ/Δ^* and three WT biological replicates, we then examined the gene expression of differentiated retinal cell types. For all cell retinal types except Müller glia, there is a reduction in the average FPKM value in R0-36*^Δ/Δ^* organoids (Fig. 4H).

During Stage 3, human retinal organoids complete maturation by forming photoreceptor outer segments (Phillips et al., 2014). To examine cell types present in mature retinal organoids, we performed single cell RNA sequencing (scRNA-seq.) on two biological replicate R0-36*^Δ/Δ^*organoids, two R3-36*^Δ/Δ^* organoids, and three WT organoids. Preliminary data suggests that R0- 36*^Δ/Δ^* retinal organoids contain an unusually high proportion of Müller glial cells (Fig. 4I).

Taken together, these data suggest that R0-36*^Δ/Δ^* organoids recapitulate microphthalmia, display a defect in proliferation, and do not produce retinal cell types in normal proportions, indicating a defect in RPC differentiation.

### There is a loss of ON cone bipolar neurons in the R3-16*^Δ/Δ^* human retinal organoids

Considering that our previous work found a complete loss of bipolar neurons in the *Vsx2* R3*^Δ/Δ^* mouse, and that this enhancer displays strong evolutionary conservation, we next wanted to assess how deletion of hR3-16 affects bipolar neurons in a model of human development (Honnell et al., 2022). The presence of all neural retinal cell types in R3-16*^Δ/Δ^* retinal organoids was first assessed by immunohistochemistry, RNA-seq, and qRT-PCR. Interestingly, each retinal cell type is present, however aberrant bipolar neuron morphology and laminar position was observed in organoids stained by G0α (Fig. 5A). Furthermore, these retinal organoids express bipolar neuron genes *GRM6, PRKCA*, and *VSX2* as determined by qRT-PCR and bulk RNA-seq (Fig. 5B, Supplemental Dataset 1). These data initially suggest that bipolar neurons are present in R3-16*^Δ/Δ^* organoids. However, considering that there was abnormal G0α staining, we decided to further investigate this phenotype by scRNA-seq.

**Figure 5.**
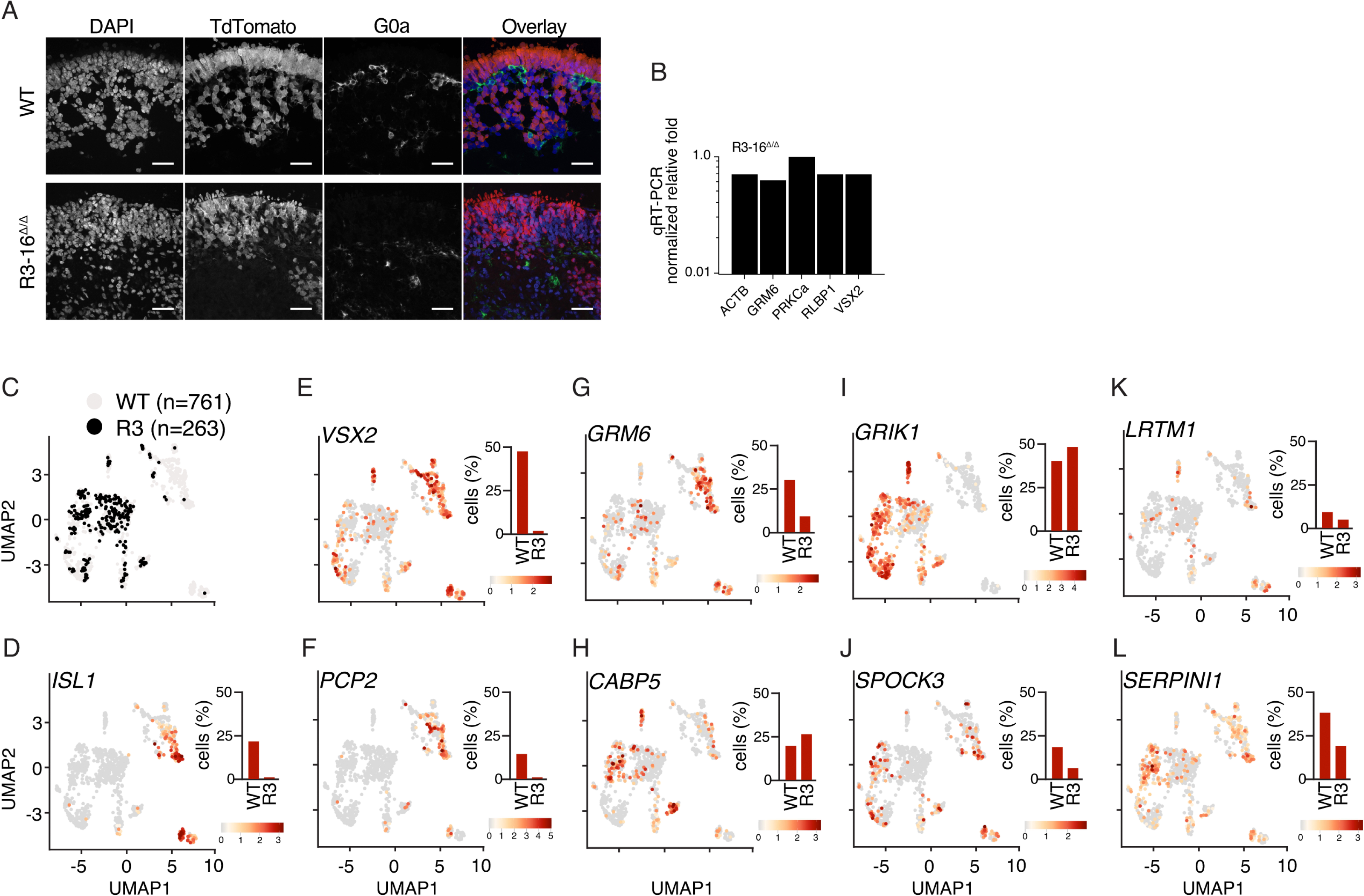
Region 3 Human retinal organoids display a loss of ON cone bipolar neurons. A. Micrograph of WT and R3-16^Δ/Δ^ retinal organoids showing immunofluorescence of DAPI (blue), TdTomato marking photoreceptors (red), and G0α marking bipolar neurons (green). Scale bar: 25 uM **B** qRT-PCR of genes expressed in bipolar neurons. **C** UMAP of Stage 3 WT (grey) and R3-16^Δ/Δ^ (black) retinal organoids. There are three WT biological replicates and two R3-16^Δ/Δ^ biological replicates. **D-L** UMAPs comprising bipolar neurons for specific genes. Heatmap displaying enrichment of gene expression (cool tones: lower expression, warm tones: high expression). Barplots displaying the proportion of cells from each sample expressing a particular gene.

Recent studies have molecularly classified bipolar neurons to identify 15 subtypes which broadly fall into the categories of ON cone bipolar, OFF cone bipolar, or rod bipolar (RB) (Shekhar et al., 2016). To examine the presence of bipolar neuron subtypes in R3-16*^Δ/Δ^* human retinal organoids, we used scRNA-seq with two R3-16*^Δ/Δ^* and three WT biological replicates.

Bipolar neurons from all five samples were isolated by computational analyses (Fig. 5C). The bipolar neurons were further categorized by established ON, OFF, or RB molecular markers (Haverkamp et al., 2003, (Shekhar et al., 2016). Interestingly, R3-16*^Δ/Δ^* bipolar neurons do not express the ON cone bipolar neuron markers *ISL1, VSX2,* and *PCP2,* and express *GRM6* at low levels compared to bipolar neurons of WT samples (Fig. 5D-G). However, both samples express RB marker, *CABP5*, and OFF cone bipolar neuron maker, *GRIK1* (Fig. 5H-1). These data suggests that deletion of hR3-16 in human retinal organoids leads to a loss of ON cone bipolar neurons. We further explored the possibility that this enhancer deletion is specific to a subtype(s) of ON cone bipolar neurons by examining distinct markers of known subtypes, however further orthogonal approaches and analyses are required (Fig. 5J-K).

## Discussion

Since their introduction a decade ago, super-enhancers have been of interest to developmental biologists because of their ability drive expression of genes encoding master regulatory transcription factors controlling cell identity and cell fate. Super-enhancers have been found to be important for normal developmental processes, and have been implicated as drivers of disease due to enrichment of disease-associated genetic variation (Hnisz et al., 2013). We believe that the importance of super-enhancer function across development is underscored by their characteristic evolutionary conservation in vertebrates (Zhang et al., 2022, (Khan et al., 2018, (Pérez-Rico et al., 2017). That is, genomic sequences within SEs have remained largely consistent over time, perhaps because they are critical for proper organogenesis and thus continued survival. Indeed, in the case of *Vsx2* we have identified distinct conserved modules within a SE that are necessary for normal retinal development in both mice and human-derived retinal organoids. Remarkably, inserting the human Region 0 module into a mouse with microphthalmia rescues eye size and vision, suggesting that this sequence is functionally interchangeable between species. To our knowledge, this is the first example of a functional interspecies enhancer rescue in vivo.

### Identification of Vsx2 SE modules

Four *Vsx2* SE regions of interest were initially identified by inspecting evolutionarily conserved sequences on the UCSC genome browser (Kent et al., 2002, (Raney et al., 2013). Boundaries were determined by visual examination and used for all plasmid and CRISPR-Cas9 knockout designs. Previous studies have identified enhancers fundamental in development by inserting conserved human sequences into mouse and examining their ability to drive reporter expression (Pennacchio et al., 2006). We used this approach with the *Vsx2* SE regions of interest to examine the ability of the human ortholog to drive reporter expression in mouse retinal cell types. Region 0 and Region 3 had cell-type specific reporter expression and the results were consistent regardless of genome build. To understand the degree of similarity between genome builds, we aligned and compared the mouse sequences to the human orthologs using the National Library of Medicine Nucleotide BLAST tool (Altschul et al., 1990, (Morgulis et al., 2008). For Region 0, BLAST identified one highly similar (89%) sequence spanning 584 bp. For Region 3, BLAST identified two sequences with that are 91% and 89% similar. Upon further analysis, these sequences are Region 3-c and Region 3-d, respectively. However, as described above, Region 3-d electroprated samples displayed more GFP+ bipolar neurons in comparison to Region 3-c (Fig S1C). Our previous ChIP-seq and scATAC data characterized Region 3 is an enhancer, however, these methods do not have the resolution to identify adjacent enhancer candidates. These results nod to the limitations of relying solely on integrated epigenetic analyses to identify enhancer elements. Furthermore, while highly conserved sequences are suggestive of regulatory regions, functional analyses are still necessary.

### Vsx2 enhancer involvement in environmental adaptation

Super-enhancers are implicated in driving normal development as well as disease (Hnisz et al., 2013, (Whyte et al., 2013, (Parker et al., 2013). However, their contribution to phenotypic diversity among species is still being explored. A previous study by the Clark Lab examined the evolutionary rate of subterranean mammalian enhancers implicated in eye development (Partha et al., 2017). This was guided by the premise that despite not being related, many subterranean mammals have poor eyesight, thus unique pressures of the underground environment may promote convergent evolution. They identified 17 eye-specific enhancers in the mole that mutated at significantly accelerated rates. Strikingly, upon mining the data, two of these enhancers are Region 0 and Region 3. We propose that the *Vsx2* SE may be a driver of phenotypic diversity in evolving species. There are other species with unusual eyes, such as the “owl” monkey (*Aotus*) which in contrast to the mole or shrew, has extremely large eyes (Dyer et al., 2009). The development of large eyes is believed to aid foraging at night, however the molecular mechanisms conferring this trait are unknown. Future studies examining evolutionary rate of super-enhancers in this species may resolve the origin of this phenotype.

### Vsx2 SE function in human retinal organoids

Retinal organoids serve as a powerful tool for modeling human retinal development and disease (Foltz and Clegg, 2019, (Kruczek and Swaroop, 2020, (Wahle et al., 2023, (Ludwig et al., 2023, (Saha et al., 2022, (Phillips et al., 2014). To understand the role of the *Vsx2* SE modules in human retinal organoids, we individually deleted Region 0 or Region 3 and assessed their growth and development over three established developmental stages (Capowski et al., 2019). One observed strength of this model is that we were able to generate hundreds of retinal organoids for each deletion line. Furthermore, the double reporter line served as an indicator of achieving a retinal lineage. However, we did observe some variation in organoid structure across biological replicates. Notably, some organoids contained a dense cellular core, while others had a hollow core. This was a limitation when scoring the proportion of apoptotic cells and instead, we scored only the outer layer which exhibited structural consistency. In this case, we cannot make definitive conclusions about the differences in overall cell death between retinal organoid lines. Instead, our interpretation is that there is no difference in cell death in the outer layer, which contains photoreceptors, bipolar neurons, amacrine cells, and horizontal cells.

Interestingly, deleting hR3-16 in human retinal organoids does not eliminate bipolar neurons as previously observed in the mouse, but instead may eliminate a sub type of bipolar cell, ON cone bipolar neurons (ON BCs). Initially, we overlooked this observation because neither IHC markers used, Vsx2 and G0α, are specific to ON BCs. However, scRNA-seq analyses later revealed a distinct absence of this cell type. Rigorous analyses are ongoing to further examine and validate this phenotype, as well as assess the subtypes of ON BCs affected. This result suggests that development of rod and OFF bipolar neurons is evolutionarily conserved, but the mechanisms by which ON BCs develop in humans is unclear.

To our knowledge, mutations in these regions have not been examined in individuals with microphthalmia or blindness. Considering that SEs harbor an increase in disease-associated risk variants, and that these regions are known to mutate at accelerated rates in low vision species, we propose that Region 0 and Region 3 could be novel drivers of microphthalmia or blindness in humans.

## Materials and Methods

### Mouse Strains

All animal procedures and protocols were approved by the St. Jude Laboratory Animal Care and Use Committee under protocol number 393-100500. All studies conform to federal and local regulatory standards. Mice were housed on ventilated racks on a standard 12 hour light-dark cycle. Wild-type C57BL/6J mice were purchased from the Jackson Laboratory (Bar Harbor, ME, #000664). For timed pregnancy, individual male mice were housed with 3 females in a single cage. Plugged/pregnant females (identified by visual examination), were isolated and embryos or pups were harvested at the appropriate time. Both males and females were combined for this study. Conserved regions within the *Vsx2* SE were identified by examination of evolutionary conservation in the UCSC mm10 genome build. Human Region 0 Enhancer mouse models were created using CRISPR-Cas9 technology.

### RNA isolation

RNA was extracted from individual TRIzol (Life Technologies) preparations via a phenol-chloroform extraction. Samples were first dissociated by pipetting retina and TRIzol vigorously. A 1:4 volume of chloroform (Sigma) was then added to each sample and incubated at room temperature for 3 min followed centrifugation at 12,000 3 g at 4C for 15 min. The aqueous layer was then transferred to a siliconized Eppendorf tube followed by the addition of 2.0 mL glycogen (Roche) and 500 mL isopropanol (Fisher Scientific). Samples were incubated at room temperature for 10 min followed by centrifugation at 12,000 3 g at 4C for 15 min. Samples were then washed twice with ice-cold 80% EtOH (Fisher) to remove salts, resuspended in DEPC H2O, and the concentration was determined with a NanoDrop (Thermoscientific).

Libraries were prepared from 500 ng total RNA with the TruSeq Stranded Total RNA Library Prep Kit according to the manufacturer’s directions (Illumina). Paired-end 100-cycle sequencing was performed on HiSeq 2000 or HiSeq 2500 sequencers according to the manufacturer’s directions (Illumina).

### qRT-PCR

cDNA was made from 200 ng of RNA from H9, JHDR, 1B6, 1C9, 5D2, 8H6, 8E10, 5B2, 1C2, 4F9 organoid lines (Applied Biosystems 4387406). cDNA was loaded onto a Custom TaqMan Array Card (Applied Biosystems 4342249) run on a QuantStudio 7 Flex (ThermoScientific) system. “Undetermined” values were set to a Ct of 40 as the limit of detection of the assay.

### RNA-seq

RNA was quantified using the Quant-iT RiboGreen assay (Life Technologies) and quality checked by 2100 Bioanalyzer RNA 6000 Nano assay (Agilent,) 4200 TapeStation High Sensitivity RNA ScreenTape assay (Agilent,) or LabChip RNA Pico Sensitivity assay (PerkinElmer) prior to library generation. Libraries were prepared from total RNA with the TruSeq Stranded Total RNA Library Prep Kit according to the manufacturer’s instructions (Illumina, PN 20020599). Libraries were analyzed for insert size distribution on a 2100 BioAnalyzer High Sensitivity kit (Agilent Technologies,) 4200 TapeStation D1000 ScreenTape assay (Agilent Technologies,) or Caliper LabChip GX DNA High Sensitivity Reagent Kit (PerkinElmer.) Libraries were quantified using the Quant-iT PicoGreen ds DNA assay (Life Technologies) or low pass sequencing with a MiSeq nano kit (Illumina) Paired end 100 cycle sequencing was performed on a NovaSeq 6000 (Illumina). For PCA analysis, only protein-coding genes with the annotation level (https://www.gencodegenes.org/pages/data_format.html) in 1 (verified loci) and 2 (manually annotated loci) were included in the analysis. With input of read counts for all samples, counts per million mapped reads (CPMs) were obtained by using the function *cpm* in the edgeR package. Genes were removed when the corresponding CPMs for all samples were smaller than the CPM whose corresponding raw read count is 10. Then, the top 3000 most variable genes were selected by ranking the mean absolute deviation (MAD) of the log_2_-transformed CPMs in descending order. Based on these most variable genes, the PCA analysis was performed on the log_2_-transformed CPMs by using the *prcomp* function available in the standard R language. The top two principal components (PCs) were used to draw the PCA figures.

### Vision Testing

The OptoMotry system from CerebralMechanics was used to measure the optomotor response of the CRISPR gene-edited mice. Briefly, a rotating cylinder covered with a vertical sine wave grating was calculated and drawn in virtual three-dimensional (3-D) space on four computer monitors facing to form a square. CRISPR gene-edited mice standing unrestrained on a platform in the center of the square tracked the grating with reflexive head and neck movements. The spatial frequency of the grating was clamped at the viewing position by repeatedly recentering the cylinder on the head. Acuity was quantified by increasing the spatial frequency of the grating until an optomotor response could not be elicited. Contrast sensitivity was measured at spatial frequencies between 0.1 and 0.45 cyc/deg. The tester was blinded to genotype until after testing was complete. Bar plot displays mean with SEM.

### GFP Reporter Assay

0.5 uL of plasmid mixture (2 ug/uL of enhancer plasmid and 0.5 ug/uL of a pCig2-H3.3- scarlet plasmid, a normalization control generously gifted to us from the Solecki Lab, resuspended in HBSS (Corning, 21-022-CV)) was co-electroporated into the sub retinal space of C57/BL6 mice at P0. Mouse retinae were harvested at P21 for GFP amplification immunostaining. Experiments were performed in biological triplicates for each enhancer plasmid. One 40X confocal image from three retina were scored for each construct. GFP+, Scarlet+ cell types were counted and divided by Scarlet+ (electroporation control) cells to calculate the percentage of GFP+ cells for each cell type. Cells were assigned to specific cell types based on their location and morphology. Bar plot displays mean and SD for each cell type for each construct.

### Organoid EdU Labeling and Scoring

EdU labeling was performed per manufacturer’s instructions (Click-iT EdU imaging kit, Invitrogen, catalog C10340), and DNA was stained with 0.2 μg/ml DAPI (Sigma-Aldrich).

Briefly, retinal organoids in individual wells containing 1 mL of 3D-RDM were given 1 uL of EdU 1-hour prior to fixing and cryoembedding. Organoids were then fixed by 4% PFA overnight, embedded by O.C.T. Compound (Scigen 4583), and cryosectioned at 10 um. Sections were fixed by 4% PFA, washed with 3% BSA-PBS, incubated with 0.5% Triton X-100 in PBS (Sigma T9284), and washed again. Cryosections were imaged using a Zeiss LSM 700 confocal microscope using a 40X lens. Three images containing at least two biological replicates were scored for each cell line. Bar plots display the mean and SD of manual scoring.

### Organoid Caspase-3 Scoring

For each genotype or human retinal organoids, 3 images from at least 2 biological replicates were collected. The fields were selected randomly using the DAPI channel in order to minimize bias in the Caspase channel. Images were collected and then total nuclei and Caspase+ nuclei were scored. Nuclear fragments were not scored. The number of Caspase + nuclei across the 3 images on a given section were combined and the total number of nuclei were combined and the ratio and percentage were calculated. The data for the two sections were averaged and the standard deviation was calculated. The individual datapoints from independent sections were plotted along with mean and SD.

### Imaging

Images were taken with the Zeiss LSM 700 confocal microscope using the 40X lens.

Brightness and contrast were modified for images presented in the figures for the IF studies. Raw original data are available for all datasets and probes.

### Statistics and Reproducibility

Mice of both sexes were randomly selected for analyses. Investigators were blinded to cell line when handling human retinal organoids. Investigators were blinded when scoring images. No statistical method was used to predetermine sample size. There were no instances in which repeat experiments yielded conflicting results, suggesting reproducibility of our experiments. GraphPad Prism 8 software was used to calculate statistical measures. No data were excluded from the analyses.

## Note: Roles of Manuscript Authors

V.H. and M.A.D. conceived and designed the study, V.H., S.S., B.T. and M.A.D. collected the data, V.H., J.L.N., and C.R. performed computational analysis, A.L. and C.B. created the hESC double reporter cell line, V.H. and M.A.D., analyzed and interpreted the data., V.H. and M.A.D. drafted the manuscript.

## Acknowledgements

We thank Shondra Pruett-Miller and Jonathon Klein of the Center of Advanced Genome Engineering and Valerie Stewart of the Neuroembryology Core for aiding with the creation of the genetically modified mouse models and cell lines, Everest Ouyang for maintaining the mouse colony and performing vision testing, Abbas Shirinifard of the Neuroimaging Analysis Lab for developing a program to measure organoid photographs, and Xitiz Chamling and Donald J. Zack for helping generate and generously sharing their transgenic hESC double reporter line. Figures were created using BioRender.com.

## Data Availability

The raw sequencing data generated in this study will be publicly available in the GEO database upon publishing. All other relevant data supporting the key findings of this study can be found within the article and its Supplementary Information files or from the corresponding author upon reasonable request.

---

**Figure S1.**
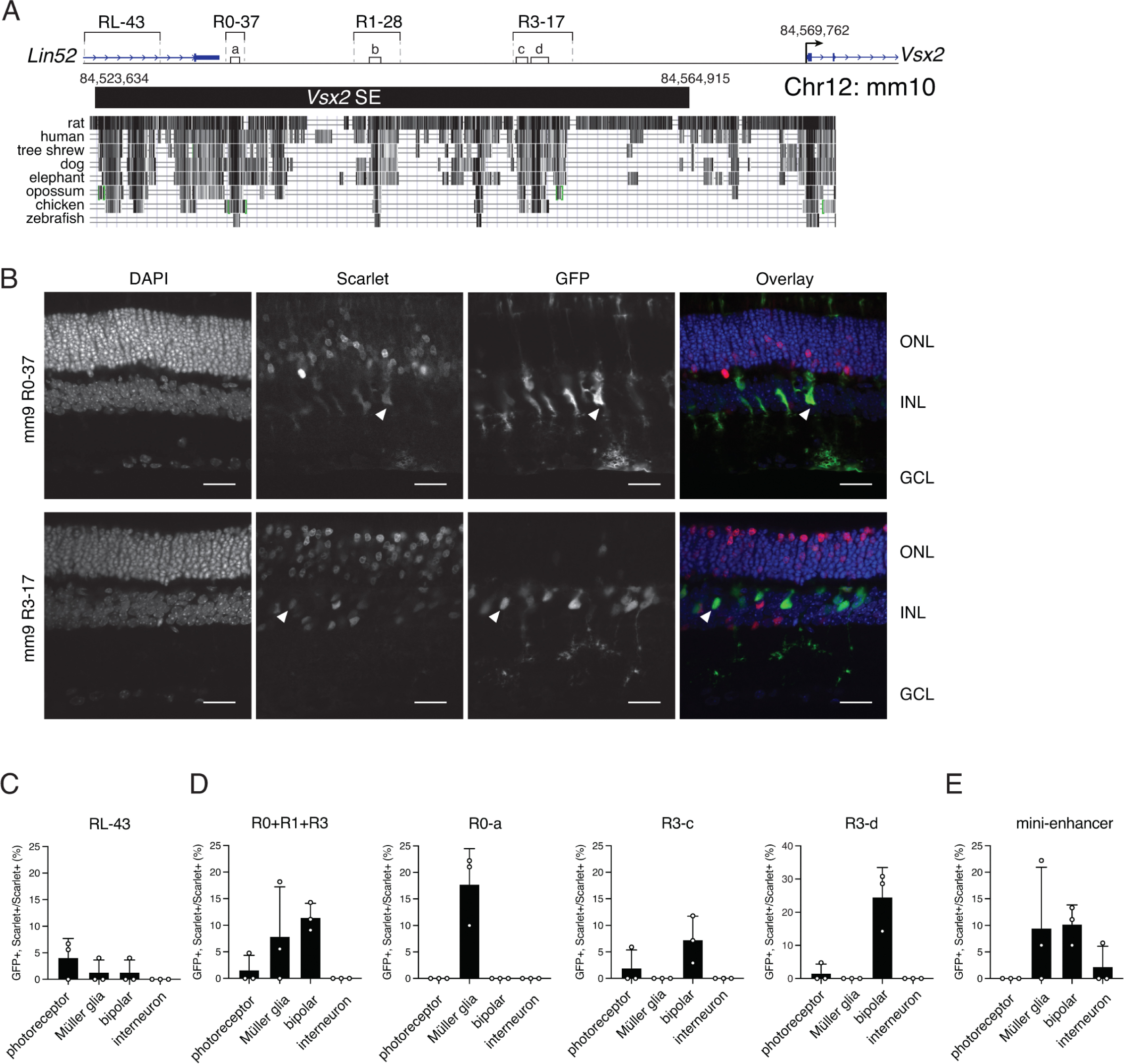
Mouse Vsx2-SE elements drive reporter expression in a cell type-specific manner. **A** Drawing of the original Vsx2 SE identified by H3K27Ac ChIP-seq (black bar) and the original mouse deletions with coordinates in mm10. Within these regions are additional refined subregions used for cloning. Evolutionarily conserved sequences across vertebrates are displayed below each Region deletion (black segments). **B** Micrographs of GFP (green) and Scarlet (red) reporter expression at P21 from square wave electroporation of P0 mice. Reporters containin mouse R0-37 and R3-17 subcloned upstream. Arrows indicate Müller glia for the R0- 37 fragment and bipolar neurons for the R3-17 fragment. Scale bar: 25 uM **C-E** Bar plot showing mean and standard deviation of three biological replicates for each mouse reporter construct. Abbreviations: RL, RegionLin52.

**Figure S2.**
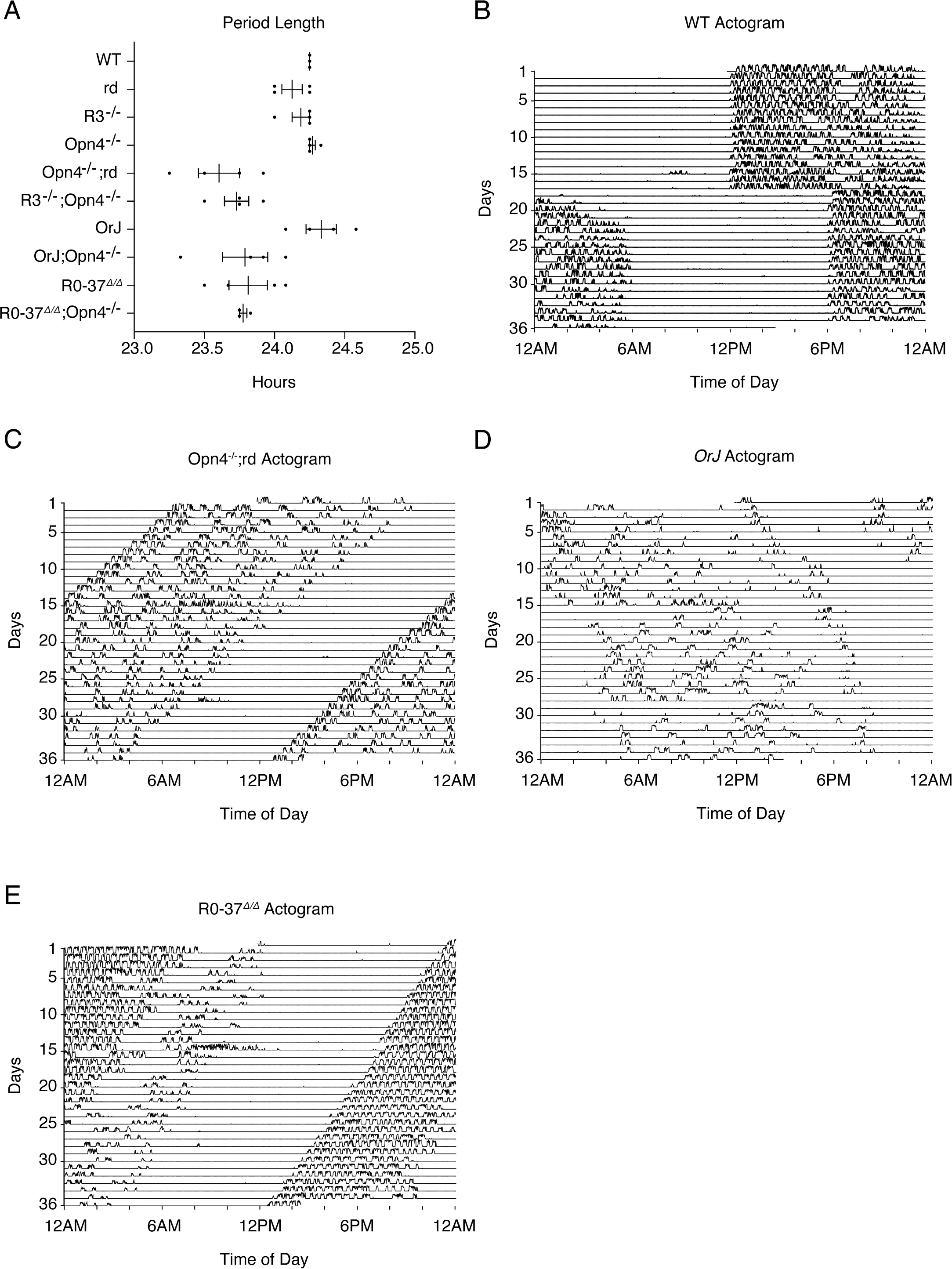
*OrJ* and R0-37^Δ/Δ^ mice do not photoentrain. **A** Period length of the 10 mouse strains tested. n= 3 biologically independent mice for each mouse strain. **B-E** Representative actograms of WT, Opn4^-/-^;rd, *OrJ*, and R0-37^Δ/Δ^ adult mice over 36 days. Lights turned off at 12PM and on at 12AM days 1-17. Lights tuned off at 6PM and on at 6AM days 18-36. Running wheel activity is displayed by peak amplitude.

**Figure S3.**
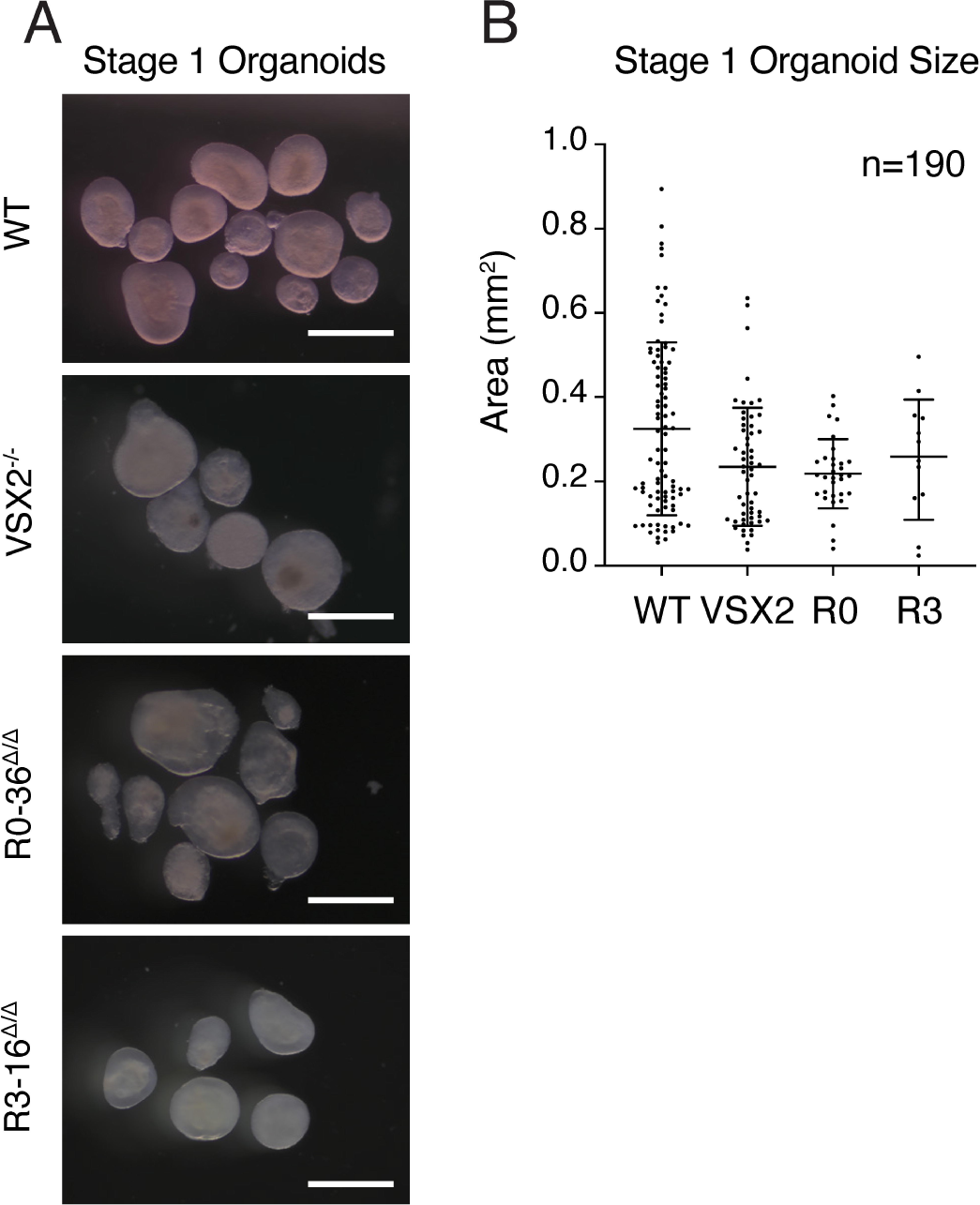
Stage 1 human retinal organoid size. **A** Stage 1 photographs of retinal organoid lines. Organoids display a phase-bright outer rim, consistent with morphological labeling at Stage 1. Scale bar: 0.5 mm. **B** Scatter dot plot of organoid area in mm^2^. Each dot represents a single retinal organoid. 190 retinal organoids were scored.

